# Pseudoalignment facilitates assignment of error-prone Ultima Genomics reads

**DOI:** 10.1101/2022.06.04.494845

**Authors:** A. Sina Booeshaghi, Lior Pachter

## Abstract

We analyze single-cell RNA-seq data sequenced with Ultima Genomics technology and find high error rates in and near homopolymers. To compensate for these errors, we explore the use of pseudoalignment for read assignment, and find that it can perform better than standard read alignment. Our pseudoalignment read assignment for Ultima Genomics data is available as part of the kallisto-bustools kb-python package available at https://github.com/pachterlab/kb_python.

## Introduction

Despite numerous improvements in DNA sequencing technology and dramatic reductions in the price of sequencing over the past fifteen years [1,2], the cost of sequencing can limit the scope of projects for biology labs [3], and is a barrier to adoption of routine sequencing in the clinic [4]. The recently unveiled Ultima Genomics sequencing technology [5] has been advertised as providing a solution to these challenges by way of delivering high-throughput sequencing at a small fraction of the cost of current technologies [6]^1^.

For single-cell RNA-seq analysis, a pilot project conducted by scientists at the Broad Institute and at Ultima Genomics found that the “data show comparable results to existing technology” [9]. However, in examining the preprint we noticed that this claim is mostly based on an apples-to-oranges comparison of (on average) 174 bp long Ultima Genomics cDNA reads to 55 bp long Illumina cDNA reads (Supplementary Figure 1). Furthermore, lower quality scores for the Ultima Genomics reads than for the Illumina reads (Figure 1) motivated us to analyze the data to see whether higher error rates in Ultima Genomics reads reduce alignment rates, and consequently degrade expression estimates for genes.

**Figure 1:**
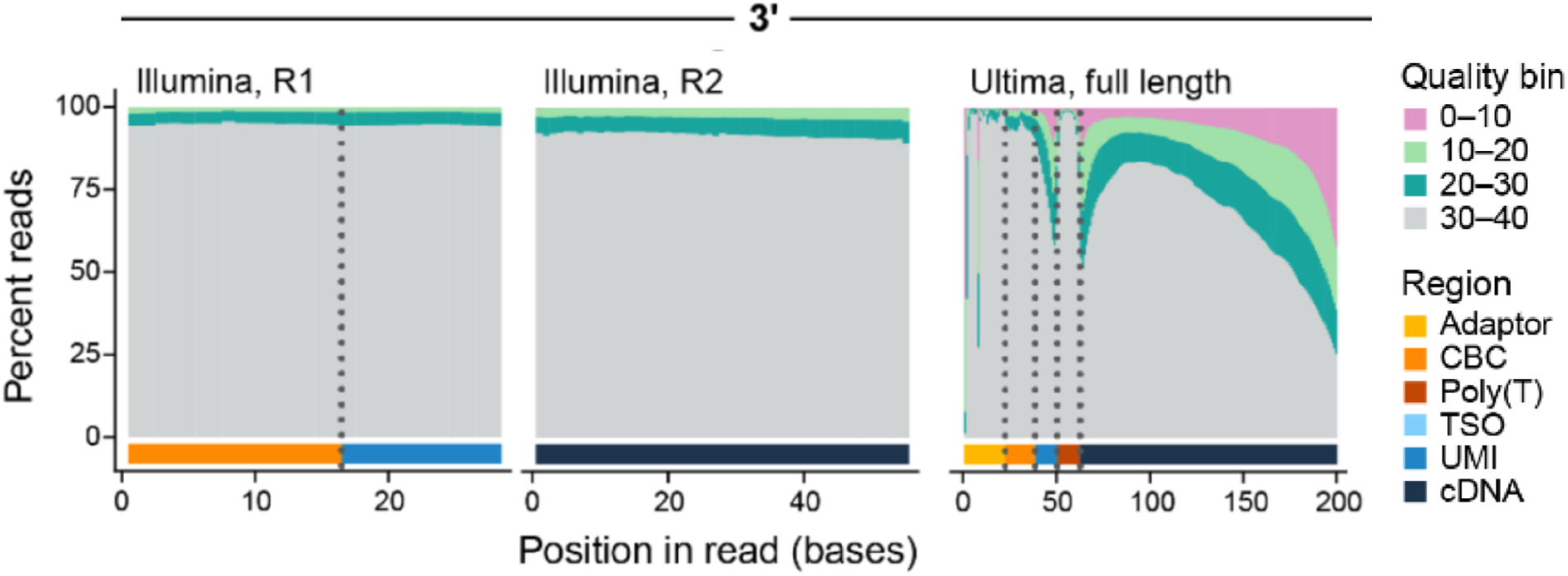
A reproduction of part of Extended Data Figure 1 from [9] which is licensed under the CC-BY 4.0 license. The Illumina reads display high quality across the barcode, UMI and cDNA sequences. The Ultima Genomics reads have lower quality at and around the Poly(T) tract, and also degraded quality after 100 bp.

## Results

[9] prepared PBMC libraries (SRX14293374) that were sequenced with both Ultima Genomics (SRR18145555) and Illumina (SRR18145553) sequencers. To perform a like-to-like comparison of (SRR18145553 and SRR18145555), we trimmed the Ultima Genomics reads to a maximum length of 55bp so as to match the length of the Illumina reads. This trimming was similar to the procedure implemented by [9] for the analysis underlying their Extended Data Figure 3H. We then pre-processed the data using kallisto-bustools [10,11], which we modified in order to be able to parse the Ultima Genomics data (Supplementary Figure 2). This resulted in gene count matrices derived from both the Ultima Genomics and Illumina sequenced cDNA libraries (see Methods). Our results corroborated the findings of [9] that at a high-level, the “data show comparable results” (Supplementary Figure 3). However, we noticed that not all genes had similar numbers of counts, and to understand why that may be the case we examined the nuclear gene with the highest difference, which was *TMSB4X*. This gene happens to be the 10th most highly expressed gene in the Illumina dataset. We found a 1.96x-fold difference in UMI counts depending on whether the library was sequenced with Ultima Genomics or Illumina; [9] also identified *TMSB4X* as an outlier in a comparison of Illumina vs. Ultima Genomics (see Simmons et al. 2022 Supplementary Table 2) but did not investigate the cause for the difference.

To understand why the Ultima Genomics UMI counts were much lower than the Illumina UMI counts, we re-aligned the reads to the *TMSB4X* gene using HISAT2 [12], and examined the alignments with the IGV browser [13]; Figure 2). The number of aligned Ultima Genomics reads was more than 4 times lower than the number of aligned Illumina reads (Table 1), and we noticed a much higher incidence of errors in the Ultima Genomics reads, specifically around homopolymer runs such as the (T)_8_ homopolymer near the 3’ end of the gene. This is consistent with the drop in quality scores near the (T)_n_ homopolymers resulting from the poly(A) tracts at the 3’ end of genes in [9]. To quantify the differences, we computed the error rates for Illumina and Ultima Genomics and found that the Ultima Genomics error rate was 10-fold higher (Table 1). We note that the errors displayed in Figure 2 and the estimates in Table 1 are only lower bounds for the error rate of Ultima Genomics because we could not align the reads with the most errors (Supplementary Figure 4). In addition to Ultima Genomics displaying a much higher mismatch error rate than Illumina for the *TMSB4X* gene, particularly concerning is the much higher rate of insertions and deletions (Figure 3a), with Ultima Genomics producing insertions and deletions up to 7 bp long. For context, liquid biopsies can benefit from accurate sequencing of 1 error per 10 million bases [14], and the accuracy of Ultima Genomics for *TMSB4X* is worse than 80,000 indels per 10 million bases. These errors are not only present at and around the (T)_8_ homopolymer in the *TMSB4X* gene, but are also apparent at and around much shorter homopolymers down to 3bp (Figure 2).

**Figure 2:**
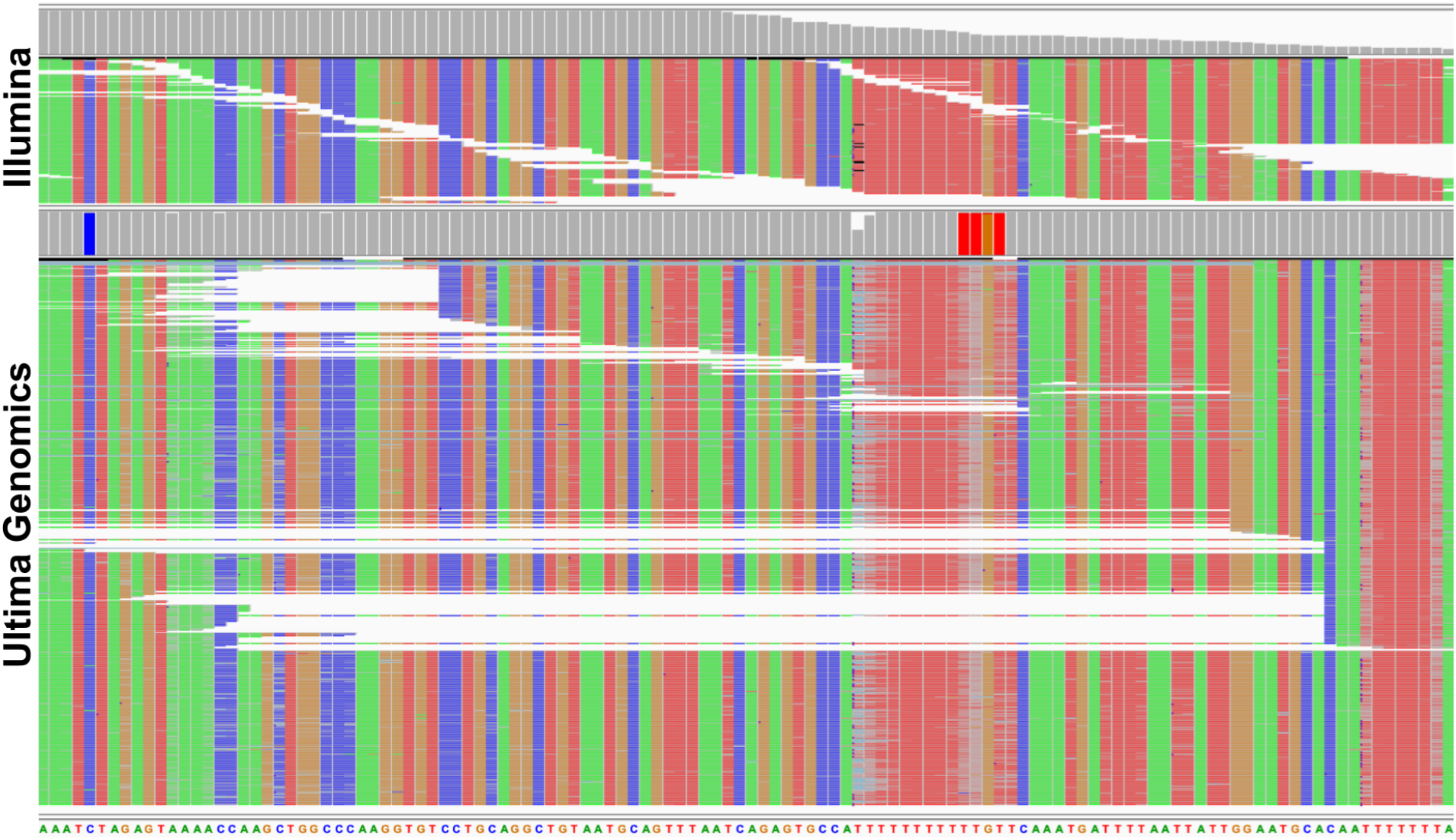
Illumina and Ultima Genomics reads aligning to the *TMSB4X* gene viewed on the IGV browser [13]. The gray streaks in the Ultima Genomics alignments reveal far more mismatches than in the Illumina alignments, especially around homopolymers. The coverage tracks above the alignments show that Ultima Genomics has a large number of indels at the (T)_8_ homopolymer whereas Illumina has far fewer indels.

**Figure 3:**
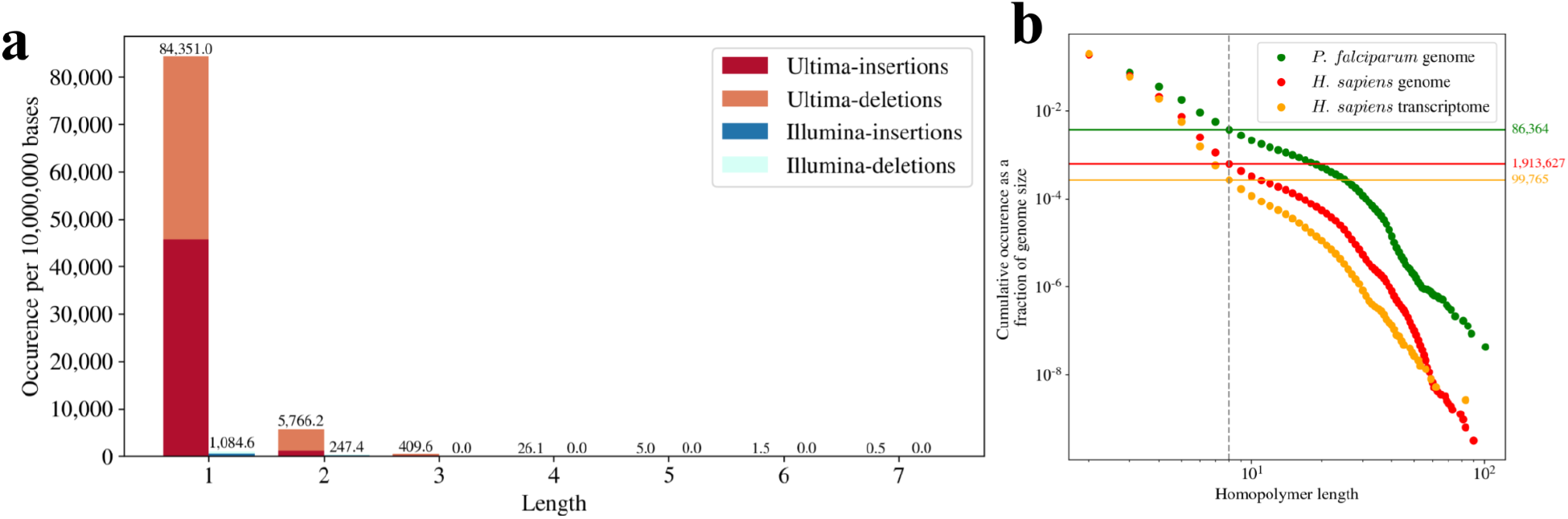
(a) The number of insertion and deletion errors in the *TMSB4X* gene for the Ultima Genomics and Illumina technologies. (b) The number of homopolymers (reported as a fraction of genome size) in the *H. sapiens* genome and transcriptome, as well as in the AT-rich *P. falciparum* genome.

**Table 1:**
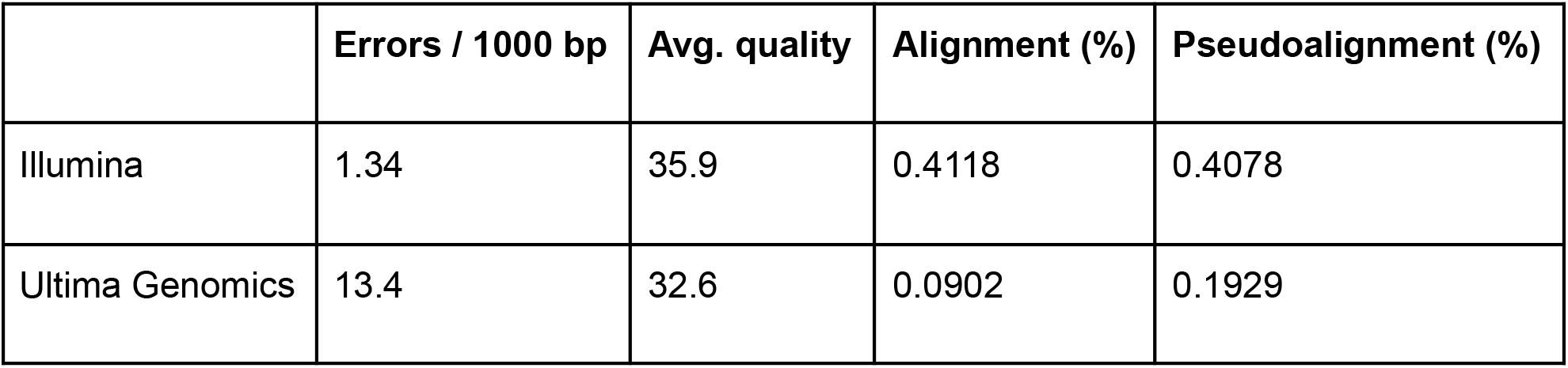
Summary *TMSB4X* gene alignment and pseudoalignment statistics for the Ultima Genomics and Illumina technologies.

In light of the overall concordance in counts between Ultima Genomics and Illumina when reads were pseudoaligned (Supplementary Figure 3), we hypothesized that pseudoalignment, which is robust to errors in reads, could rescue some of the unaligned reads originating from the *TMSB4X* gene. We found that 0.1929% of Ultima Genomics reads pseudoaligned, which is 2.14 times higher than the fraction of aligned reads (0.0902%). In the case of Illumina reads, the higher base-call quality translated to little difference between the fraction of aligned (0.4118%) and pseudoaligned (0.4078%) reads. Thus, for *TMSB4X*, pseudoalignment recovered more than double the number of reads that could be assigned, demonstrating that it is a method well-suited to counting of error-prone Ultima Genomics reads for single-cell RNA-seq. Nevertheless, the Ultima Genomics error rate is so high that even pseudoalignment fails to rescue all the reads.

## Discussion

In the Ultima Genomics preprint [5], the company discusses the challenges of sequencing homopolymers with their non-terminating chemistry, but touts an accuracy of 90% for homopolymers of length 8, and characterizes this as “good accuracy up to length 8-10 bases”. We find that in the Ultima Genomics single-cell RNA-seq reads, the homopolymer challenge presents as more acute than what may be imagined from summary statistics. The *TMSB4X* example demonstrates that Ultima Genomics displays poor performance not only in regions with long homopolymer runs, extreme GC content, or highly repetitive sequences. We found that the Ultima Genomics difficulties in sequencing *TMSB4X* transcripts in PBMCs are also present in the K562 cell line; the data in [15] shows a 3.9 fold reduction in Ultima Genomics alignments vs Illumina alignments.

While the genome-wide accuracy of Ultima Genomics technology is likely to be better than for the *TMSB4X* gene, a comprehensive assessment of the error profiles was not possible because at the time of writing of this preprint, the data in [5] has not been released (the preprint states that “the data will be made available in the near future”). However, regardless of the genome-wide performance of Ultima Genomics technology, it is worth noting that improvement of sequencing technology is best guided by understanding worst-case performance.

Furthermore, the poor performance of Ultima Genomics on the *TMSB4X* gene is clinically relevant, because *TMSB4X* is possibly a biomarker for renal cancer as it has been shown to be predictive for survival [16–18]. Moreover, *TMSB4X* is relevant for several clinical applications [19].

While pseudoalignment of Ultima Genomics reads provides an improvement over standard alignment due to robustness to error, the improvement may not suffice for all genes, resulting in biases that may be difficult to compensate for during analysis. This problem may be addressable by improved alignment or pseudoalignment algorithms that are robust to homopolymers in the reference sequence. Some algorithmic ideas for this challenge have been developed in the context of other sequencing technologies that have poor performance in sequencing homopolymers; see, for example, [20] that was developed for Ion Torrent sequencing technology.

Unfortunately, in its current form, it seems that while Ultima Genomics sequencing may be promising in the long run, it is not currently a direct replacement for Illumina or BGI sequencers, both of which have much higher accuracy. The problems we found in sequencing regions with homopolymers, will be magnified in whole-genome sequencing applications. We counted the number of homopolymers in the human genome and found 1,913,627 homopolymers of at least length 8; the number of homopolymers can be even higher than that in other genomes (Figure 3b). Even for single-cell RNA-seq, the homopolymer problem has implications beyond just read assignment. [9] observed that the poly(T) homopolymers in their reads corrupted the UMIs, and they therefore had to truncate the last 3 bases of each UMI (implemented by replacing the last 3 bases of each UMI with AAA). While this compensates for mismatches due to the poly(T) homopolymers, it has drawbacks in terms of accurate molecule counting [10].

Hopefully Ultima Genomics will eventually find a way to reduce error rates so they can be competitive with existing technologies. The “many degrees of freedom” provided by their architecture design [5] is possibly reason for optimism. The genomics community is already benefiting from cost reductions following the introduction of BGI sequencers [21–23], and will benefit even further if Ultima Genomics can similarly reduce sequencing costs.

## Methods

The code to reproduce all the figures and results in the preprint is available at https://github.com/pachterlab/BP_2022 and provides a complete description of the methods.

## Data Availability

The Illumina single-cell data is available on GEO under accession GSM6297378 and the Ultima single-cell data is available on GEO under accession GSM6297379. The Illumina Perturb-seq data is available on GEO under accession GSM6190598 and the Ultima Perturb-seq data is available on GEO under accession GSM6190599. Previously the data had been made available on GEO under accession GSM5917802 but the authors of [9] had to modify the original GEO entry because the Ultima data was being misidentified as Illumina data.

## Software

- anndata 0.7.6 [24]
- bustools 0.40.0 [10,25]
- IGV 2.13.0 [13]
- kallisto 0.48.0 [11]
- kb-python 0.27.2 [10]
- ffq 0.2.1 [26]
- gget 0.1.1 [27]
- HISAT2 2.2.1 [12]
- htslib 1.10 [28]
- Matplotlib 3.5.1 [29]
- Numpy 1.21.6 [30]
- Pandas 1.3.5 [31]
- samtools 1.10 [32]
- seqkit 0.12.0 [33]

## Footnotes

1 The “cost of sequencing” is not a well-defined concept. It can refer to only the cost of reagents for a sequencing run, or it can include other costs such as library preparation, personnel, analysis, space etc. Thus, statements such as “the $100 genome” are not meaningful; for example the oft cited NIH website for cost estimates [7] includes numerous production costs other than just instruments and reagents. This is because the reagent costs can be dominated by other costs [3]. Furthermore, there is no accepted accuracy or completeness standard for the “sequence of a human genome”. For example Nebula Genomics sells a $99 genome [8], but at 0.4x coverage rather than the more common 30x.

## References

1. LeMieux J. All Aboard The Genome Express. Genetic Engineering & Biotechnology News. Mary Ann Liebert, Inc., publishers; 39:34, 35, 38, 40–12019;

2. Goodwin S, McPherson JD, McCombie WR. Coming of age: ten years of next-generation sequencing technologies. Nat Rev Genet. 17:333–512016;

3. Li H, Wu K, Ruan C, Pan J, Wang Y, Long H. Cost-reduction strategies in massive genomics experiments. Marine Life Science & Technology. 1:15–212019;

4. Schwarze K, Buchanan J, Fermont JM, Dreau H, Tilley MW, Taylor JM, et al.. The complete costs of genome sequencing: a microcosting study in cancer and rare diseases from a single center in the United Kingdom. Genet Med. 22:85–942020;

5. Almogy G, Pratt M, Oberstrass F, Lee L, Mazur D, Beckett N, et al.. Cost-efficient whole genome-sequencing using novel mostly natural sequencing-by-synthesis chemistry and open fluidics platform. bioRxiv.

6. : Ultima Genomics Delivers the $100 Genome. https://www.ultimagenomics.com/blog/ultima-genomics-delivers-usd100-genome (2022). Accessed 2022 Jun 3.

7. Kris A. Wetterstrand MS: DNA Sequencing Costs: Data. Genome.gov. NHGRI; https://www.genome.gov/about-genomics/fact-sheets/DNA-Sequencing-Costs-Data (2019). Accessed 2022 Jun 4.

8. : Nebula Genomics - 30x Whole Genome Sequencing - DNA Testing. https://web.archive.org/web/20220525222324/https://nebula.org/whole-genome-sequencing-dna-test/ (2022). Accessed 2022 Jun 4.

9. Simmons SK, Lithwick-Yanai G, Adiconis X, Oberstrass F, Iremadze N, Geiger-Schuller K, et al.. Single cell RNA-seq by mostly-natural sequencing by synthesis. bioRxiv.

10. Melsted P, Booeshaghi AS, Liu L, Gao F, Lu L, Min KHJ, et al.. Modular, efficient and constant-memory single-cell RNA-seq preprocessing. Nat Biotechnol. 39:813–82021;

11. Bray NL, Pimentel H, Melsted P, Pachter L. Near-optimal probabilistic RNA-seq quantification. Nat Biotechnol. 34:525–72016;

12. Kim D, Paggi JM, Park C, Bennett C, Salzberg SL. Graph-based genome alignment and genotyping with HISAT2 and HISAT-genotype. Nat Biotechnol. 37:907–152019;

13. Robinson JT, Thorvaldsdóttir H, Winckler W, Guttman M, Lander ES, Getz G, et al.. Integrative genomics viewer. Nat Biotechnol. 29:24–62011;

14. Higgins J, Pratt G, Valentine CC, Williams LN, Salk JJ. Redefining “Gold Standard”: Ultra-Sensitive Characterization of Commercial DNA Standards with Duplex Sequencing. Blood. 134:20932019;

15. Replogle JM, Saunders RA, Pogson AN, Hussmann JA, Lenail A, Guna A, et al.. Mapping information-rich genotype-phenotype landscapes with genome-scale Perturb-seq. bioRxiv.

16. Morita T, Hayashi K ’ichiro. Tumor Progression Is Mediated by Thymosin-β4 through a TGFβ/MRTF Signaling Axis. Mol Cancer Res. 16:880–932018;

17. : Expression of TMSB4X in renal cancer - The Human Protein Atlas. https://www.proteinatlas.org/ENSG00000205542-TMSB4X/pathology/renal+cancer/KIRC (2018). Accessed 2022 Jun 3.

18. Uhlen M, Zhang C, Lee S, Sjöstedt E, Fagerberg L, Bidkhori G, et al.. A pathology atlas of the human cancer transcriptome. Science. 2017; doi: 10.1126/science.aan2507.

19. Crockford D, Turjman N, Allan C, Angel J. Thymosin beta4: structure, function, and biological properties supporting current and future clinical applications. Ann N Y Acad Sci. 1194:179–892010;

20. Feng W, Zhao S, Xue D, Song F, Li Z, Chen D, et al.. Improving alignment accuracy on homopolymer regions for semiconductor-based sequencing technologies. BMC Genomics. 17 Suppl 7:5212016;

21. Drmanac S, Callow M, Chen L, Zhou P, Eckhardt L, Xu C, et al.. CoolMPS™: Advanced massively parallel sequencing using antibodies specific to each natural nucleobase. bioRxiv.

22. Hahn O, Fehlmann T, Zhang H, Munson CN, Vest RT, Borcherding A, et al.. CoolMPS for robust sequencing of single-nuclear RNAs captured by droplet-based method. Nucleic Acids Res. 49:e112021;

23. LeMieux J: MGI Delivers the $100 Genome at AGBT Conference. Genetic Engineering and Biotechnology News. https://www.genengnews.com/news/mgi-delivers-the-100-genome-at-agbt-conference/ (2020). Accessed 2022 Jun 4.

24. Virshup I, Rybakov S, Theis FJ, Angerer P, Alexander Wolf F. anndata: Annotated data. bioRxiv.

25. Melsted P, Ntranos V, Pachter L. The barcode, UMI, set format and BUStools. Bioinformatics. academic.oup.com; 35:4472–32019;

26. Gálvez-Merchán Á, Min KH (joseph), Pachter L, Sina Booeshaghi A. Metadata retrieval from sequence databases with ffq. bioRxiv.

27. Luebbert L, Pachter L. Efficient querying of genomic reference databases with gget. bioRxiv.

28. Bonfield JK, Marshall J, Danecek P, Li H, Ohan V, Whitwham A, et al.. HTSlib: C library for reading/writing high-throughput sequencing data. Gigascience. 2021; doi: 10.1093/gigascience/giab007.

29. Hunter JD. Matplotlib: A 2D Graphics Environment. Comput Sci Eng. Institute of Electrical and Electronics Engineers (IEEE); 9:90–52007;

30. Harris CR, Millman KJ, van der Walt SJ, Gommers R, Virtanen P, Cournapeau D, et al.. Array programming with NumPy. Nature. 585:357–622020;

31. Mckinney W: Pandas: A foundational Python library for data analysis and statistics. https://www.dlr.de/sc/portaldata/15/resources/dokumente/pyhpc2011/submissions/pyhpc2011_submission_9.pdf (2011). Accessed 2022 Jun 5.

32. Li H, Handsaker B, Wysoker A, Fennell T, Ruan J, Homer N, et al.. The Sequence Alignment/Map format and SAMtools. Bioinformatics. 25:2078–92009;

33. Shen W, Le S, Li Y, Hu F. SeqKit: A Cross-Platform and Ultrafast Toolkit for FASTA/Q File Manipulation. PLoS One. 11:e01639622016;

