## Supplementary Material for "Pseudoalignment facilitates assignment of error-prone Ultima Genomics reads"

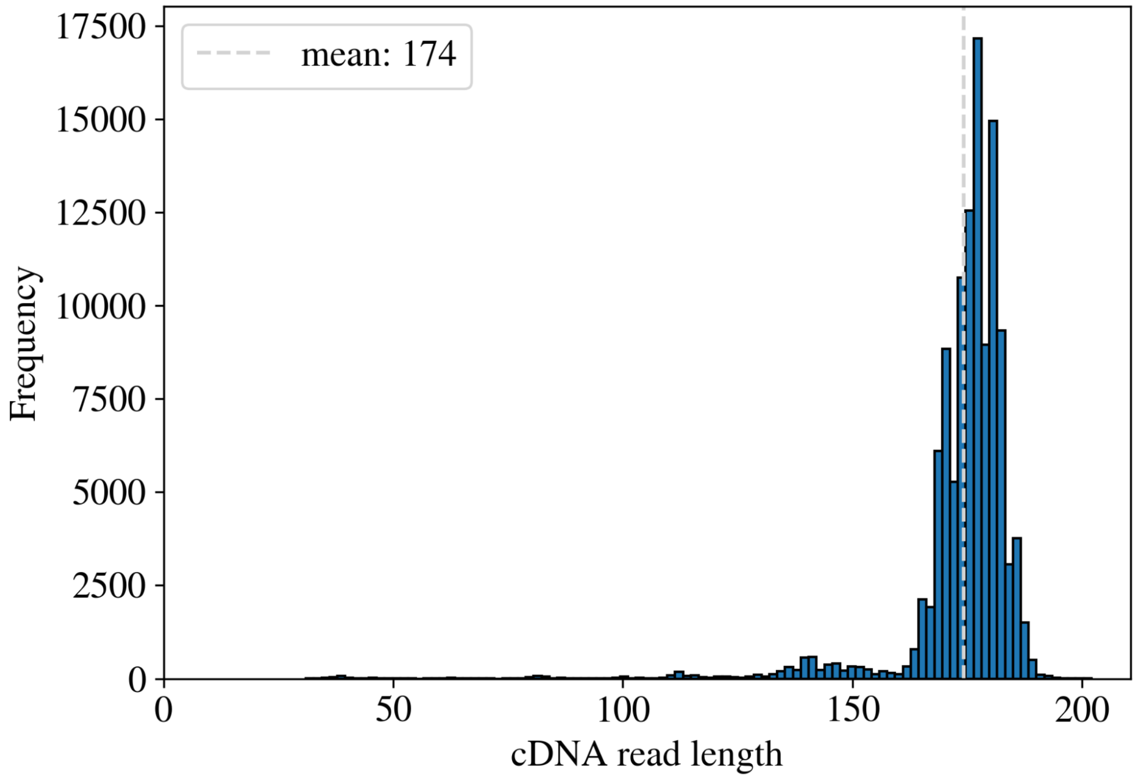


**Supplementary Figure 1:** Distribution of Ultima Genomics cDNA read lengths.

```
$ kb count -x 10xv3_Ultima -o out -i index.idx -g t2g.txt read1.fastq.gz
```

**Supplementary Figure 2**: Pre-processing Ultima Genomics single-cell RNA-seq with kallisto-bustools.

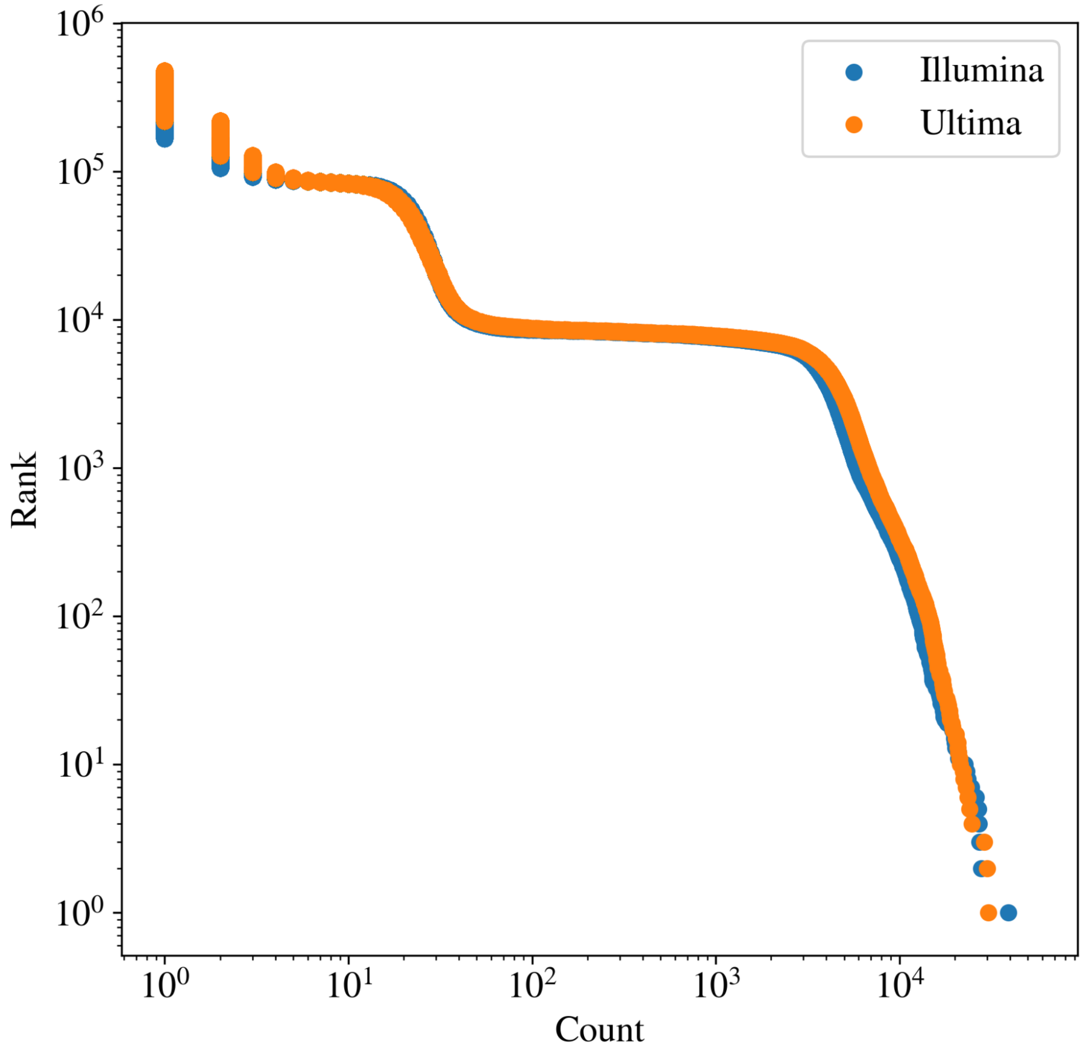


**Supplementary Figure 3**: Knee plots for the Ultima Genomics (55bp CDNA reads) - Illumina (55bp cDNA reads) comparison.

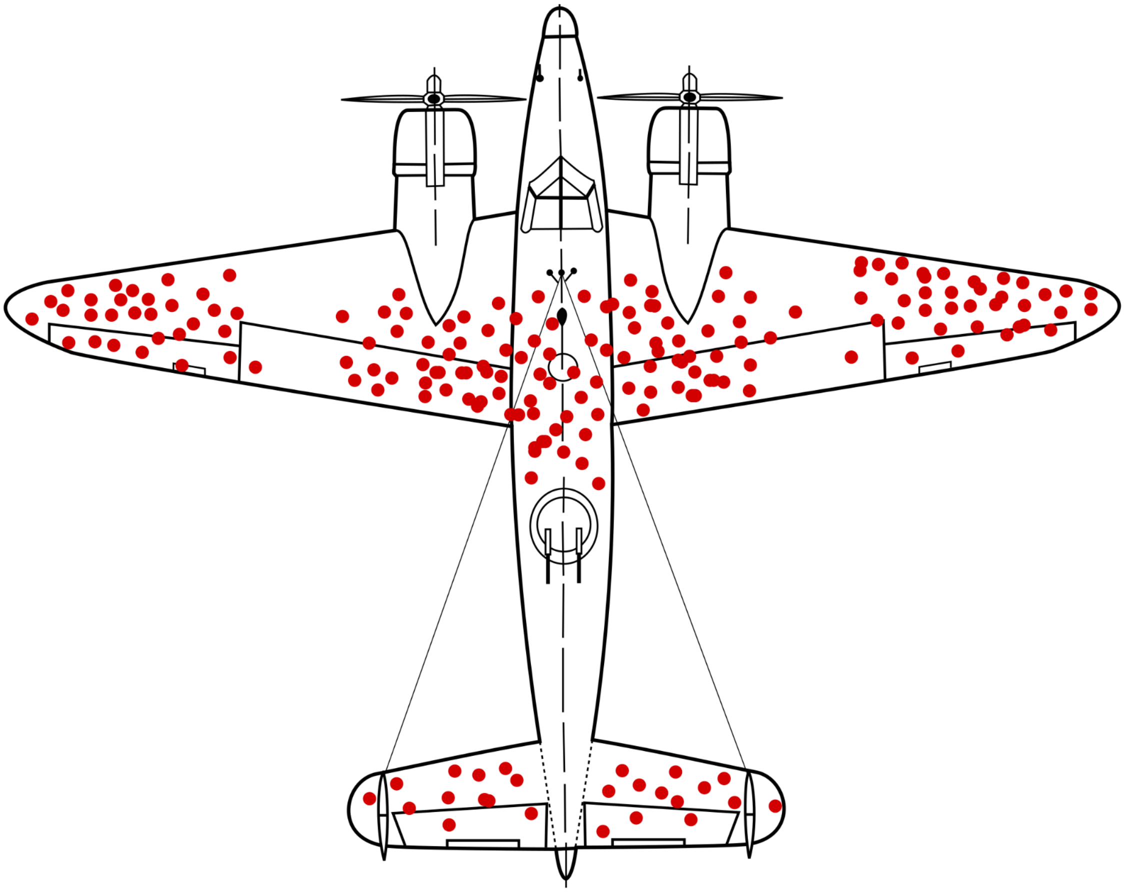

**Supplementary Figure 4**: An illustration of survivorship bias in the work of Abraham Wald on "A Method of Estimating Plane Vulnerability Based on Damage of Survivors" during WWII (1940). Source of image: Wikipedia (license CC BY-SA 4.0).
